# Individual Differences in Delay Discounting are Associated with Dorsal Prefrontal Cortex Connectivity in Youth

**DOI:** 10.1101/2023.01.25.525577

**Authors:** Kahini Mehta, Adam Pines, Azeez Adebimpe, Bart Larsen, Dani S. Bassett, Monica E. Calkins, Erica Baller, Martin Gell, Lauren M. Patrick, Raquel E. Gur, Ruben C. Gur, David R. Roalf, Daniel Romer, Daniel H. Wolf, Joseph W. Kable, Theodore D. Satterthwaite

## Abstract

Delay discounting is a measure of impulsive choice relevant in adolescence as it predicts many real-life outcomes, including substance use disorders, obesity, and academic achievement. However, the functional networks underlying individual differences in delay discounting during youth remain incompletely described. Here we investigate the association between multivariate patterns of functional connectivity and individual differences in impulsive choice in a large sample of youth. A total of 293 youth (9-23 years) completed a delay discounting task and underwent resting-state fMRI at 3T. A connectome-wide analysis using multivariate distance-based matrix regression was used to examine whole-brain relationships between delay discounting and functional connectivity was then performed. These analyses revealed that individual differences in delay discounting were associated with patterns of connectivity emanating from the left dorsal prefrontal cortex, a hub of the default mode network. Delay discounting was associated with greater functional connectivity between the dorsal prefrontal cortex and other parts of the default mode network, and reduced connectivity with regions in the dorsal and ventral attention networks. These results suggest that delay discounting in youth is associated with individual differences in relationships both within the default mode network and between the default mode and networks involved in attentional and cognitive control.

## INTRODUCTION

Delay discounting (DD) is a measure of impulsive decision-making (Madden et al., 2003) that refers to preference for a smaller reward sooner rather than a larger reward later (Bickel et al., 2012; Epstein et al., 2010). DD predicts many real-life outcomes, such as academic achievement and social functioning (Hirsh et al., 2008; Mahalingam et al., 2016). Additionally, DD is considered an important transdiagnostic behavior that is altered across multiple clinical disorders that are characterized by impulsive decisions, including substance misuse, schizophrenia, and attention-deficit hyperactivity disorder (ADHD; Amlung et al., 2019; Chase et al., 2011; Weller et al., 2014; Ortiz et al., 2015; Lempert et al., 2019). A better understanding of the mechanisms of DD could thus inform decisions regarding early interventions for certain disorders, particularly in at-risk adolescents. However, studies that link functional brain networks defined using functional connectivity (FC) to DD in youth remain sparse. Here, we sought to understand how DD is related to individual differences in functional brain networks in a large sample of youth.

Many studies have used task-based fMRI to uncover the brain regions engaged during DD, especially key regions involved in reward valuation such as the ventral striatum and hubs of the default mode network (DMN) such as the ventromedial prefrontal cortex (vmPFC) and posterior cingulate cortex (PCC; Schüller et al., 2019; Souther et al., 2022; Kable and Glimcher, 2007; Peters and Büchel, 2010). A related but distinct approach links DD to FC at rest instead of task-based responses. Work using FC is motivated in part by behavioral data that has suggested DD is a stable trait that varies among individuals and is heritable (Kirby, 2009). FC has previously proven predictive of individual personality traits and has also been used successfully to identify neural correlates of DD (Kable and Levy, 2015). Studies of individual differences in FC related to DD often use a network-based framework, which is supported by prior research suggesting that DD relies upon interactions among multiple brain networks (Chen et al., 2017). Specifically, prior work in adults has linked impulsive choice during DD to individual differences in connectivity in regions involved in reward and valuation such as the striatum, vmPFC and PCC (Kable and Levy, 2015; Li et al., 2013; Calluso et al., 2015). Work in adults has also found that connectivity between the DMN and cognitive control networks such as the ventral attention and cingulo-opercualr networks is predictive of delay discounting – increased functional connectivity between the two typically anticorrelated networks could disrupt cognitive control and impact decisions on DD tasks (Chen, Guo, Suo, and Feng, 2018). This is consistent with the idea of a role for top-down attentional/cognitive control in delay of gratification, as indicated in previous work (Hare et al., 2014; Mischel et al., 1989).

While there have been fewer studies of children and adolescents, prior work investigating ADHD has also related individual differences in DD to connectivity in regions important for valuation, such as the nucleus accumbens (Costa Dias et al., 2013). Similarly, work in both typically developing populations and children with ADHD indicates that cognitive control regions such as the dorsolateral prefrontal cortex (dlPFC) are related to DD in adolescents (Wang et al., 2017; Rosch et al., 2018). However, results from prior work in adolescents are for the most part heterogeneous, which may be driven by two factors. First, many studies of DD and functional networks in pediatric samples have been small, increasing the risk of type I error and reducing the likelihood of replicable results (Marek et al., 2022; Button et al., 2013). Second, many studies have related DD to FC among a specific set of regions or limited set of networks, rather than evaluating the complete functional connectome. As DD is a complex cognitive process that engages multiple brain networks, such studies may not capture important differences in connectivity that are distributed across the cortex (Chen et al., 2017).

Accordingly, here we investigated how individual differences in DD are associated with connectome-wide differences in patterns of FC during adolescence. We capitalized on a large sample of 293 youth imaged as part of the Philadelphia Neurodevelopmental Cohort (Satterthwaite et al., 2014; Satterthwaite et al., 2016) who completed a DD task and resting-state fMRI. We conducted a connectome-wide association study (CWAS) to reveal DD-associated differences in the multivariate pattern of connectivity at each location in the brain (Shehzad et al., 2014; Sharma et al., 2017). While CWAS is a data-driven approach, we sought to test the hypothesis that individual differences in DD would be linked to connectivity in regions of both the DMN and networks involved in attentional control.

## METHODS

### Participants

This study considers participants who completed both neuroimaging and a DD task as part of the Philadelphia Neurodevelopmental Cohort (PNC; Satterthwaite et al., 2014); this sample largely overlaps with a previous report linking DD to individual differences in brain structure (Pehlivanova et al., 2018). Of the 1601 youths who completed neuroimaging as part of the PNC, 453 participants completed a behavioral DD task outside of neuroimaging sessions and were thus eligible for further analyses. Of these, n=2 did not meet the quality control criteria for behavioral data (Pehlivanova et al., 2018; see *DD task* section). Further, 21 participants were excluded for the following reasons: health conditions that could impact brain structure (n=19), scanning performed 12 months from the time of DD testing (n=1), and missing imaging data (n=1). An additional 137 participants were excluded due to poor quality scans, as described in the *Image quality assurance* section. Thus, a total of 293 participants (ages 9-23 years; *M =*17.18 years, *SD* = 3.10 years; 156 females, 137 males) formed the sample for our analyses after quality control.

### Ethics

This study received approval from the institutional review boards at the University of Pennsylvania and Children’s Hospital of Philadelphia. All adult participants provided informed consent. For minors, parent or guardians provided informed consent and the minor provided assent.

### DD task

The DD task consisted of 34 self-paced questions where the participant chose between a smaller amount of money available immediately or a larger amount available after a delay (Senecal et al., 2012; Pehlivanova et al., 2018). The smaller, immediate rewards ranged from $10 to $34, and the larger, delayed rewards were fixed at $25, $30, or $35 with equal frequency. Delays ranged from 1 to 171 days. Trials and task procedures were identical in content and order for all participants. The DD task was administered as part of an hour-long web-based battery of neurocognitive tests as part of a procedure used previously, on a separate day from the imaging session (Gur et al., 2010; Gur et al., 2012).

Discount rates from the DD task were calculated with hyperbolic discounting model of the form:

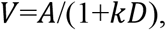

where *V* is the subjective value of the delayed reward, *A* is the amount of the delayed reward, *D* is the delay in days, and *k* is the subject-specific discount rate (Mazur, 1987; Kable and Glimcher, 2010). As in previous work, the *fmincon* optimization algorithm in MATLAB (MathWorks) was used to estimate the best fitting *k* from each participant’s choice data, assuming that choices were a logistic function of *V*s (Senecal et al., 2012; Pehlivanova et al., 2018). A higher *k* value indicates steeper discounting of delayed rewards and thus more impulsive choices. As the distribution of estimated *k* parameters is right-skewed, we applied a log-transform (log(*k*)) before further analysis.

Quality assurance of DD data was conducted as described previously: each participant’s responses were fit using a logistic regression model, with predictors including the immediate amount, delayed amount, delay, their respective squared terms, and two-way interaction terms (Pehlivanova et al., 2018). The goodness of fit of this model was assessed using the coefficient of discrimination (Tjur et al., 2009); a value less than 0.20 indicated nearly random choices and resulted in exclusion (Pehlivanova et al., 2018). As prior (Pehlivanova et al., 2018), we evaluated assocations between log(*k*) and demographic variables using a linear model.

### Image acquisition

All MRI scans were acquired using the same 3T Siemens (Erlangen, Germany) Tim Trio wholebody scanner and 32-channel head coil at the Hospital of the University of Pennsylvania. Image acquisition protocols are described in detail in previous work (Satterthwaite et al., 2014). Briefly, the magnetization-prepared, rapid acquisition gradient-echo T1-weighted (MPRAGE) image was acquired with the following parameters: TR = 1810 ms; TE = 3.5 ms; TI = 1100 ms, FOV = 180 × 240 mm^2^, matrix = 192 × 256, effective voxel resolution = 0.938 × 0.938 × 1 mm^3^. Restingstate fMRI scans were acquired with a single-shot, interleaved multi-slice, gradient-echo, echo planar imaging (GE-EPI) sequence sensitive to BOLD contrast with the following parameters: TR = 3000 ms; TE = 32 ms; flip angle = 90°; FOV = 192 × 192 mm^2^; matrix = 64 × 64; 46 slices; slice thickness/gap = 3/0 mm, effective voxel resolution = 3.0 × 3.0 × 3.0 mm^3^. Resting-state scans consisted of 124 volumes. In addition, a B0 field map was derived for application of distortion correction procedures, using a double-echo, gradient-recalled echo (GRE) sequence: TR = 1000 ms; TE_1_ = 2.69 ms; TE_2_ = 5.27 ms; 44 slices; slice thickness/gap = 4/0 mm; FOV = 240 mm; effective voxel resolution = 3.75×3.75×4 mm^3^.

### Image processing

Before the processing of both structural and functional data, a custom adolescent template was created with Advanced Normalization Tools (ANTs; Avants and Gee, 2004; Avants et al., 2011a). The template was created to minimize registration bias and maximize sensitivity to detect regional effects that can be impacted by registration error (Avants et al., 2011a). Structural images were then processed and registered to this template using the ANTs cortical thickness pipeline (Tustison et al., 2014). This procedure includes brain extraction, N4 bias field correction (Tustison et al., 2010), Atropos probabilistic tissue segmentation (Avants et al., 2011b), and the SyN diffeomorphic registration method (Avants et al., 2008; Klein et al., 2009).

The fMRI data were processed with an empirically validated preprocessing pipeline implemented in the eXtensible Connectivity Pipeline (XCP) Engine (Ciric et al., 2018). Resting-state time series preprocessing included correction of distortion induced by magnetic field inhomogeneity using FMRIB Software Library (FSL)’s FUGUE utility (Jenkinson, 2003), realignment of all volumes to a selected reference volume using MCFLIRT (Jenkinson et al., 2002), interpolation of intensity outliers in each voxel’s time series using AFNI’s 3dDespike utility and demeaning and removal of first- and second-order trends. After the despiking and detrending, the functional data were denoised using a 36-parameter confound regression model that has been shown to minimize associations with motion artifact and other nuisance variables (Ciric et al., 2017). Specifically, the confound regression model included the six framewise estimates of motion, the mean signal extracted from eroded white matter and cerebrospinal fluid compartments, the global signal, the derivatives of each of these nine parameters, and quadratic terms of each of the nine parameters as well as their derivatives. To avoid frequency mismatch, both the BOLD-weighted time series and the confound regressor timeseries were temporally filtered simultaneously using a first-order Butterworth filter with a passband between 0.01 and 0.08 Hz (Hallquist et al., 2013). Confound regression was performed using AFNI’s 3dTproject. Denoised functional images were coregistered to the T1 image using boundary-based registration (Greve and Fischl, 2009) and aligned to the study-specific adolescent template using the ANTs transform for the T1 image as above. Functional images were resampled to 4 mm^3^ isotropic voxels in the template space before CWAS for computational feasibility (Shehzad et al., 2014). However, higher spatial resolution images (2 mm^3^) were used for follow-up seed-based analyses. Throughout, all transformations were concatenated so that only one interpolation was performed.

### Image quality assurance

Some participants were excluded due to inadequate structural image quality (n=3), as determined by three expert raters (Rosen et al., 2017). As described in prior work (Satterthwaite et al., 2013; Ciric et al., 2018), a participant’s resting-state fMRI data was excluded if the mean relative root mean square (RMS) framewise displacement was higher than 0.2 mm, or if it had more than 20 frames with motion exceeding 0.25 mm (n=133). One participant was also excluded when manual inspection revealed fewer data points than expected in the resting-state scan (n=1). Our final sample thus included 293 participants. Additionally, to account for residual motion in the data that passed quality assurance, we included RMS framewise displacement as a covariate in all models.

### Multivariate distance-based matrix regression (MDMR)

We performed a CWAS using MDMR as described in detail in previous studies (Shehzad et al., 2014; Satterthwaite et al., 2015; Sharma et al., 2017). A schematic of the CWAS procedure is depicted in **Figure 1**. First, the preprocessed participant BOLD time series were used to conduct seed-based connectivity analyses at each voxel within gray matter. Specifically, Pearson’s correlation coefficient between each voxel’s time series and the time series of every other gray matter voxel (**Figure 1A &B**) was used to generate subject-level connectivity maps. Second, we summarized individual differences in FC maps by computing a distance matrix (also using Pearson’s correlation) between the connectivity matrices for every possible pairing of participants (**Figure 1C**). Third, MDMR (**Figure 1D**) was used to test how well our phenotypic variable, log(*k*), explained variation in the distances between connectivity matrices across participants. This approach provided a measure of how FC patterns across participants were impacted by individual differences in log(*k*), while controlling for the effects of age, sex (assigned at birth), and in-scanner motion (Shehzad et al., 2014; Satterthwaite et al., 2015). MDMR yields a pseudo-F statistic for each voxel, whose significance was assessed using 5,000 iterations of a permutation test to generate a null distribution. The ultimate product of this procedure was a voxel-wise *Z*-statistic map describing the association between log(*k*) and the global pattern of connectivity for each voxel (**Figure 1E**). Aligning with current recommendations to minimize false positives, the type I error rate across voxels was controlled using cluster correction with a voxel height of z > 3.09 and utilized a cluster-extent probability threshold *p* < 0.05 (Eklund et al., 2016). We also ran an analysis to explore interactions with age and sex (log(*k*)*age or log(*k*)*sex); these models included the same covariates listed above.

**Figure 1.**
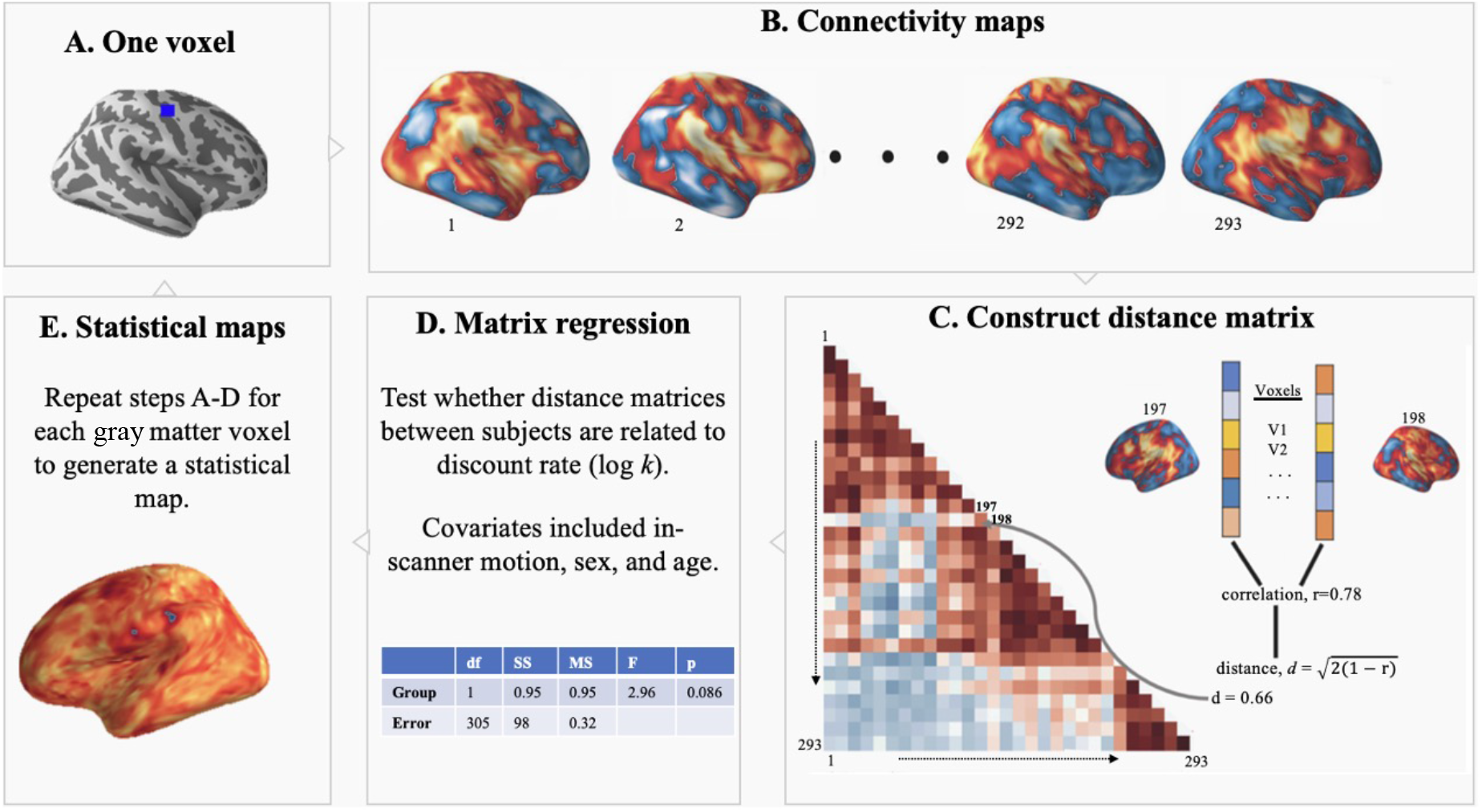
Connectome-wide analysis approach. For each gray matter voxel (**A**), a connectivity map was created for each subject (**B**), and the maps were compared in a pairwise manner (using correlation) to create a distance matrix (**C**). Multivariate distance-based matrix regression (MDMR) was used to evaluate how the multivariate patterns of connectivity encoded by these distance matrices were associated with individual differences in delay discounting while controlling for age, sex, and in-scanner motion (**D**). Permutation testing yielded a pseudo-*F* statistic and a corresponding *p* value. This procedure was repeated for each gray matter voxel, yielding a voxel-wise statistical map (**E**).

### Seed-based analyses

MDMR identified clusters where the overall multivariate pattern of connectivity is dimensionally related to DD, but it did not describe the specific pairwise FC patterns that drove the multivariate results. To characterize the direction of the effects, as in previous studies (Satterthwaite et al., 2015; Sharma et al., 2017), we conducted post-hoc seed-based descriptive analyses for each cluster returned by MDMR. Group-level seed analysis included age, sex, and in-scanner motion as covariates and was computed using a general linear model (implemented in FSL’s *flameo;* Woolrich et al., 2004). These follow-up analyses were descriptive, as the seeds were selected based on the significance of the MDMR result.

### Network enrichment testing

Given that neural activity differs across functional networks (Raut et al., 2020), we attempted to localize effects of interest within specific brain networks. Specifically, we examined whether associations with log(*k*) revealed by the seed-based analyses described above were located within one of the seven canonical large-scale brain networks (Yeo et al., 2011) using a conservative network enrichment testing procedure (see Baller et al., 2022 for details). To account for the different size of each network and the spatial autocorrelation of brain maps, statistical testing used a conservative spin-based spatial permutation procedure (Alexander-Bloch et al., 2018). Areas with positive and negative associations were evaluated separately. Briefly, statistical maps from the seed-based analysis were thresholded at |z|≽3.09 and projected onto a spherical representation of the cortical surface. This sphere was rotated 1,000 times per hemisphere to create a null distribution. For both the real and permuted data, we evaluated proportion of vertices that overlapped with each of the seven canonical functional networks. Networks were considered to have significant enrichment if the test statistic in the observed data was in the top 5% of the null distribution derived from permuted data.

### Sensitivity analyses of socioeconomic status

To probe whether our results could be driven by individual differences in socioeconomic status (SES), mean parental education was included as a model covariate in addition to age, sex, and head motion.

## RESULTS

### Connectome-wide analyses identify a region of connectivity related to DD

We sought to determine whether and how individual differences in DD were associated with complex, multivariate patterns of FC in a large sample of youth. Notably, we found no significant correlation between log(*k*) and either age or sex. However, as these variables may be strongly associated with functional connectivity, they were included as model covariates in the connectome-wide analysis.

Our connectome-wide analysis using MDMR revealed that DD was related to a multivariate pattern of FC in the left dorsal prefrontal cortex (dPFC; cluster center of gravity: x=30.9, y=43.8, z=30.3; *k*=12 voxels, *p*=1.03×10^-4^; **Figure 2**). This finding suggested a pivotal role of the dPFC, a hub of the DMN (Alves et al., 2019; Andrews-Hanna et al., 2010), in DD-related activity. We additionally evaluated models that included interactions between DD and both sex and age; the interaction effects in these models were not statistically significant.

**Figure 2.**
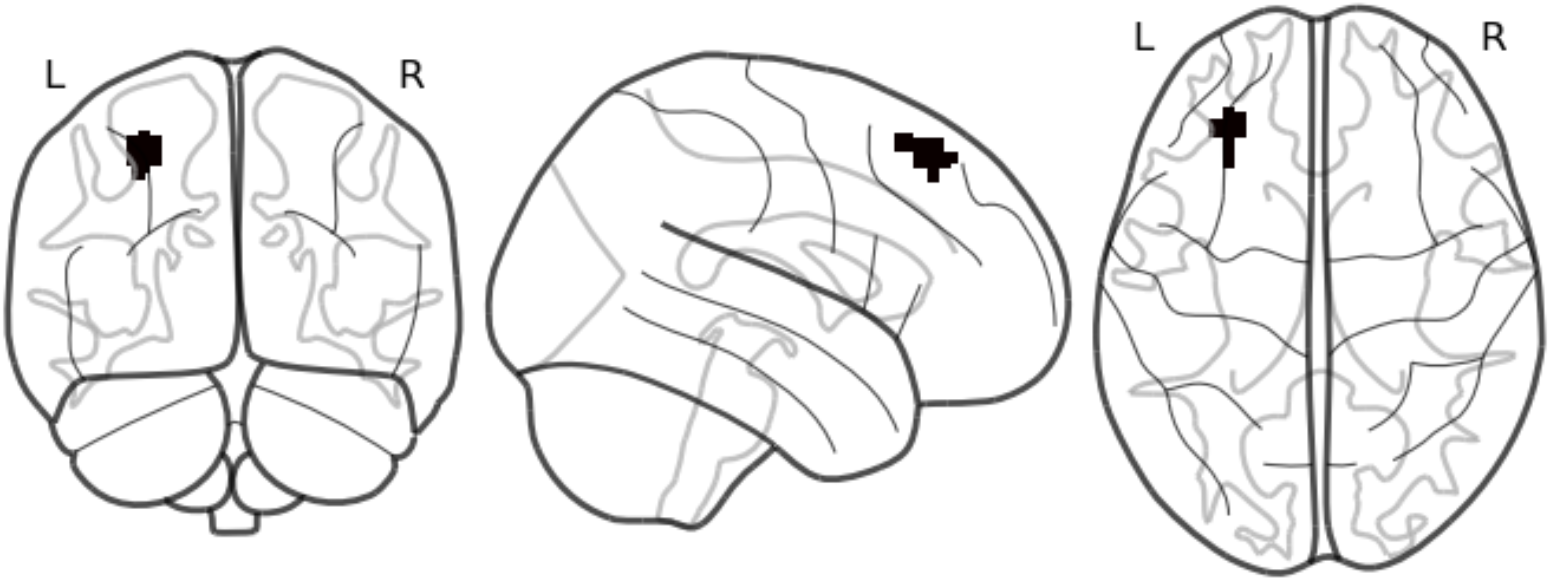
Connectome-wide analyses reveal that multivariate patterns of connectivity in the dorsal prefrontal cortex is associated with delay discounting. Volumetric depiction of the dorsal prefrontal cortex cluster identified by multivariate distance-based matrix regression. This dorsal prefrontal cortex cluster survived correction for multiple comparisons at *z*>3.09, *p*<0.05.

### DD is related to individual differences in connectivity between attentional control and default mode networks

Having localized multivariate connectivity patterns associated with DD to the dPFC, we next sought to understand the individual differences in FC associated with DD that drove the observed MDMR results. We conducted seed-based connectivity analysis using the dPFC cluster identified by MDMR. In this general linear model, we included age, sex, and in-scanner motion as covariates.

We first evaluated the mean pattern of connectivity for the dPFC cluster. Across the entire sample, the left dPFC seed was strongly connected to elements of the DMN, including the PCC and vmPFC. The seed was anticorrelated mainly to regions within the dorsal attention network (DAN), such as the inferior parietal lobule, and regions in the ventral attention network (VAN) such as the temporoparietal junction (**Figure 3)**. This connectivity profile suggests that the dPFC cluster was primarily affiliated with the DMN.

**Figure 3.**
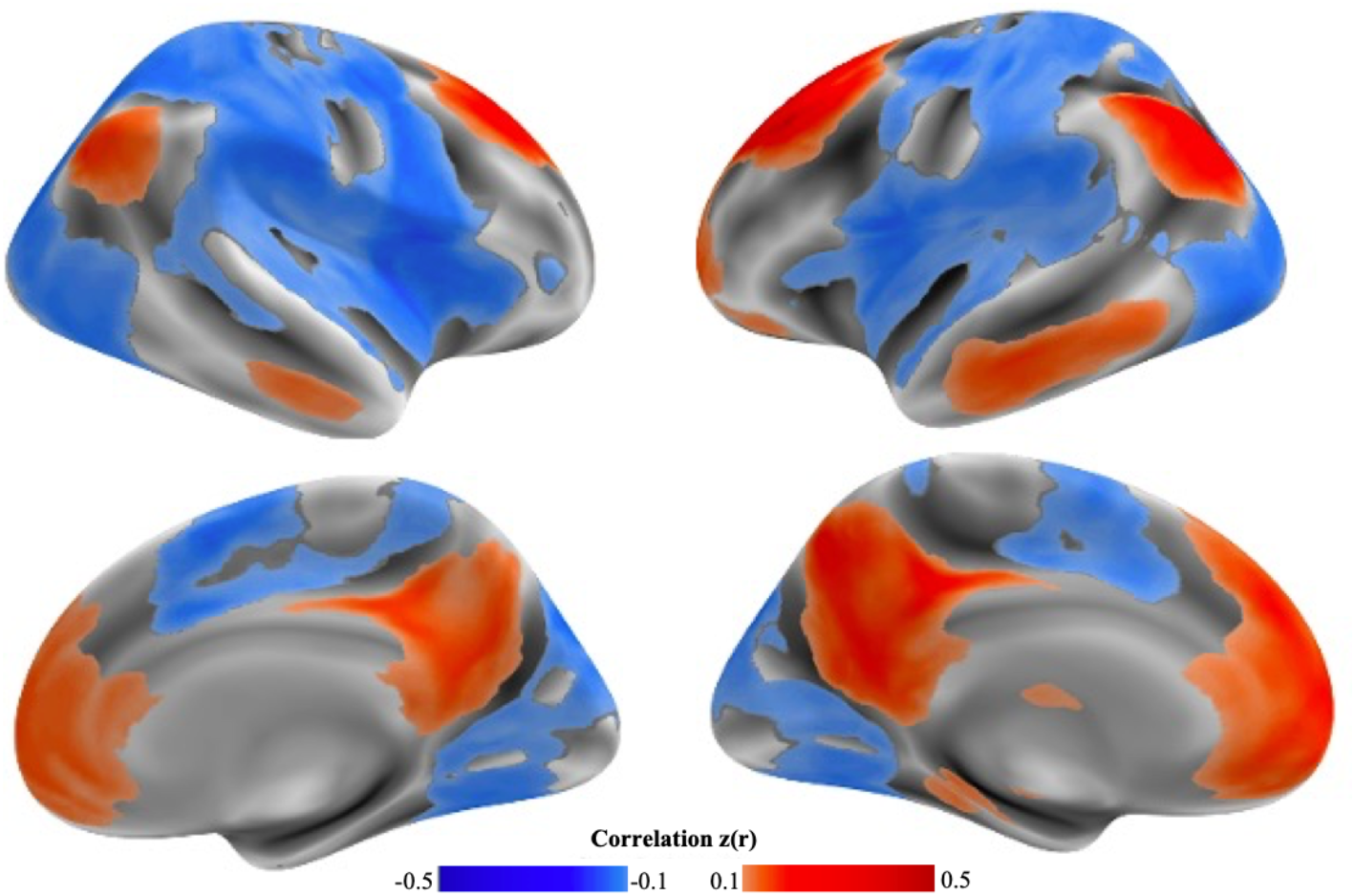
Mean connectivity of the dorsal prefrontal cortex cluster. The cluster identified by the connectome-wide association study (see Figure 2) was used as a seed to understand the connectivity profiles of the regions related to delay discounting. The left dorsal prefrontal cortex cluster had robust connectivity to other elements of the default mode network and was anticorrelated primarily with the dorsal and ventral attention network regions.

Next, we sought to determine how DD was associated with individual differences in FC from the dPFC seed identified by the connectome-wide analysis. Analysis of the cluster within the left dPFC revealed that higher rates of DD were correlated with increased connectivity between the dPFC and other elements of the DMN, including the PCC and lateral temporal cortex (**Figure 4**). In contrast, higher levels of DD were correlated with lower connectivity between the dPFC and regions within the VAN (including the temporoparietal junction and parts of the ventral frontal cortex) and the DAN (including the inferior parietal lobule and angular gyrus).

**Figure 4.**
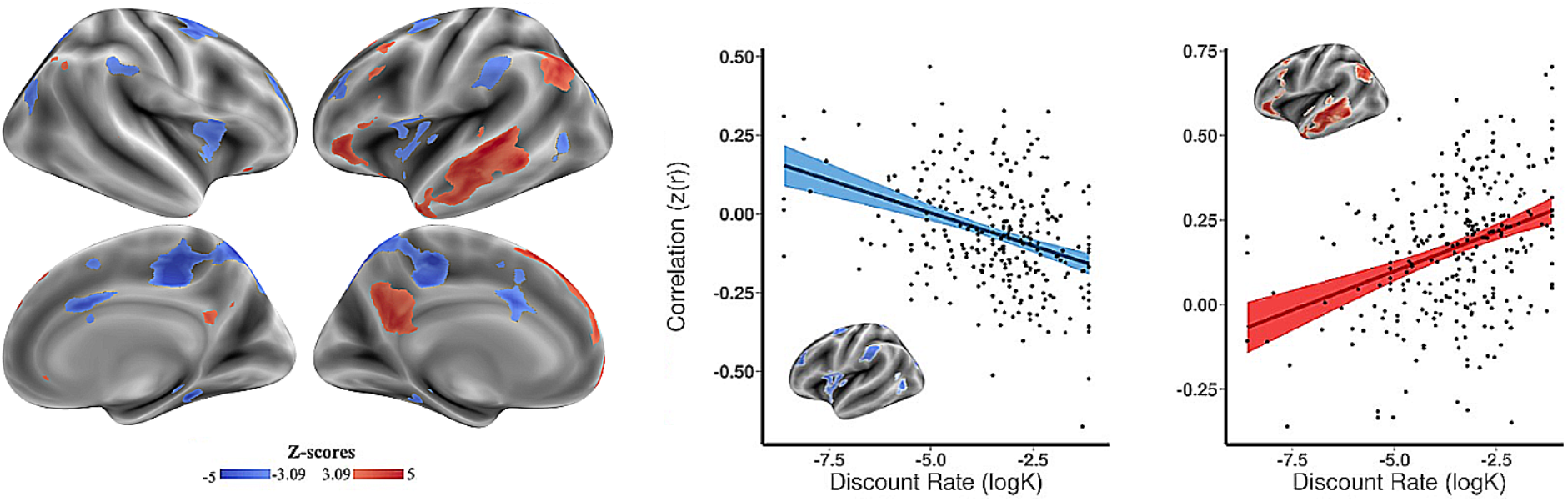
Individual differences in delay discounting are associated with dorsal prefrontal cortex connectivity to the default mode network, as well as control and attention networks. Follow-up seed-based analyses from the dorsal prefrontal cortex revealed that increased discount rate was associated primarily with increased connectivity with other elements of the default mode network (red), as well as diminished connectivity with mainly the dorsal and ventral attention network regions (blue). Maps represent patterns that drove the connectome-wide association study result rather than independent statistical tests. The maps are thresholded for display at |*z|*>3.09, p<0.05.

We next used spin-based network enrichment testing to statistically evaluate xthe spatial distribution of these effects. Enrichment testing revealed an enrichment of positive associations with log(*k*) in the DMN (*p*=0.01). In contrast, there was enrichment of negative associations in the DAN (*p*=0.02) and VAN (*p*=1.5x 10^-3^). Together, these results could suggest that DD in youth is associated with individual differences connectivity within the DMN and between the DMN and attention networks. Specifically, higher rates of discounting (more impulsive choices) are associated with greater connectivity between the dPFC and other DMN regions, but lower connectivity between the dPFC and attentional control regions.

### Sensitivity analyses

The dPFC cluster remained significant when SES was added as a model covariate as part of sensitivity analyses.

## DISCUSSION

In this study, we used a data-driven approach to identify multivariate FC patterns that underlie DD in a large sample of children, adolescents, and young adults. Our approach revealed that connectivity patterns of a region within the DMN—the dPFC—related to individual differences in DD. Further analyses revealed that higher DD was associated with increased FC of the dPFC with other regions within the DMN, and reduced FC with regions within the DAN and VAN. Taken together, these findings suggest that the dPFC may be a key node important for individual differences in impulsive choice within large-scale functional networks during resting-state imaging.

Notably, different parts of the dPFC are related to different aspects of DD, including the dlPFC, an executive control region, (Hare et al., 2014) and the dMPFC, implicated in processing future rewards and delay time (Wang et al., 2021). Other regions of the DMN, including the vmPFC and PCC, have been implicated in subjective valuation processes critical for decision-making (Pfeifer and Berkman, 2018; Bartra, McGuire, and Kable, 2013; Kable and Glimcher, 2007). Greater impulsivity has been associated with changes in how these regions represent reward features and value differences during DD tasks (Vanyukov et al., 2015; Koban et al., 2021). When interpreted in the context of these previous findings, our results suggest that stronger integration between the dPFC and other hubs of the DMN involved in valuation could be associated with impulsive decision-making, consistent with at least one smaller study in younger adults (Jung et al., 2021).

We found that participants with a greater discount rate also showed greater anticorrelation between the dPFC and networks involved in attentional and cognitive control such as the DAN and VAN. This pattern mirrors resting-state functional segregation—or increased anticorrelation between disparate brain networks (Fair et al., 2007). Our results parallel previous studies showing that resting-state functional segregation between large-scale networks—for example, the connectivity of the DMN with the cingulo-opercular network, which is involved in cognitive control (Sadaghiani, and D’Esposito, 2014)—can predict DD (Chen, Guo, Suo, and Feng, 2018). This pattern of anticorrelation with the attention networks aligns with the frequently observed dissociation between task-positive attention networks and the task-negative DMN (Fox et al., 2005). These findings suggest that stronger connections between regions within the DMN, together with weaker connections between the DMN and attentional control networks, could underlie higher DD through changes in attentional control and reward valuation.

Prior work suggests that adolescence is an important period for the general organization of large-scale FC, in which changes in connectivity adhere to a sensorimotor-association gradient that culminates in the DMN, a network clearly implicated in our analyses (Sydnor et al., 2022; Margulies et al., 2016). Further, the FC of the dPFC is known to evolve throughout adolescence with a shift from general to more differentiated abilities (Li et al., 2022). There is also evidence for functional separation between regions of the dPFC including the dmPFC and dlPFC, both regions involved in DD (Li et al., 2022). Nonetheless, we found no associations between age and DD in our work.

### Limitations

Several limitations of this work should be noted. First, our study is cross-sectional rather than longitudinal, which may have limited our ability to find associations between age and DD. However, the null effects seen in our sample do align with the inconsistent and small age effects in the DD literature as noted before (Romer, et al., 2017). Nonetheless, it is important to acknowledge that developmental changes in DD may occur at earlier ages than those studied here, and that longitudinal studies might detect developmental effects through measurement of within-person change documented in previous studies (Anandakumar et al., 2018; Achterberg et al., 2016). Second, the MDMR approach has limited sensitivity in many settings, potentially increasing the risk of type II error. For example, MDMR analysis is often insensitive to more focal changes because it summarizes differences in distributed multivariate patterns of connectivity (Misaki et al., 2018). Finally, our task used hypothetical rather than real rewards as part of the DD paradigm. However, previous research has not revealed differences between performance on DD tasks with real versus hypothetical rewards (Bickel et al., 2009).

### Conclusions

We found that the pattern of dPFC connectivity is related to individual differences in impulsive choice during youth. Multivariate patterns associated with impulsive choice were driven primarily by increased connectivity between the dPFC and other parts of the DMN, as well as diminished connectivity with attention networks. Moving forward, the results from this data-driven analysis will be important to replicate. While speculative, these results also suggest that the dPFC may be a potential target for TMS and neuromodulatory therapies for conditions where impulsivity is prominent.

## ACKNOWLEDGEMENTS

This work was supported by the U.S. National Institute of Mental Health grants MH120174 (David R. Roalf), MH119185 (David R. Roalf), R01MH117014 (Ruben C. Gur, Raquel E. Gur), R01MH119219 (Ruben C. Gur, Raquel E. Gur), R01MH120482 (Theodore D. Satterthwaite), R01MH113550 (Theodore D. Satterthwaite), R01EB022573 (Theodore D. Satterthwaite), R37MH125829, (Theodore D. Satterthwaite), RC2MH089983 (Raquel E. Gur), RC2MH089924 (Raquel E. Gur), K99MH127293 (Bart Larsen), 2T32MH019112-29A1 (Erica Baller), R01MH113565 (Daniel H. Wolf), F31MH123063-01 (Adam Pines) and Deutsche Forschungsgemeinschaft (DFG, German Research Foundation) grant 269953372/GRK2150 (Martin Gell). There are no competing interests at this time.

## DATA AVAILABILITY

The Philadelphia Neurodevelopmental Cohort is a publicly available dataset at https://www.ncbi.nlm.nih.gov/projects/gap/cgi-bin/study.cgi?study_id=phs000607.v3.p2.

## CODE AVAILABILITY

All code and a reproducibility guide for this project is available at https://github.com/PennLINC/pncitc.

